# Natural variation in *C. elegans* short tandem repeats

**DOI:** 10.1101/2022.06.25.497600

**Authors:** Gaotian Zhang, Ye Wang, Erik C. Andersen

## Abstract

Short tandem repeats (STRs) represent an important class of genetic variation that can contribute to phenotypic differences. Although millions of single nucleotide variants (SNVs) and short indels have been identified among wild *Caenorhabditis elegans* strains, the natural diversity in STRs remains unknown. Here, we characterized the distribution of 31,991 STRs with motif lengths of 1-6 bp in the reference genome of *C. elegans*. Of these STRs, 27,636 harbored polymorphisms across 540 wild strains and only 9,691 polymorphic STRs (pSTRs) had complete genotype data for more than 90% of the strains. Compared to the reference genome, the pSTRs showed more contraction than expansion. We found that STRs with different motif lengths were enriched in different genomic features, among which coding regions showed the lowest STR diversity and constrained STR mutations. STR diversity also showed similar genetic divergence and selection signatures among wild strains as in previous studies using single-nucleotide variants. We further identified STR variation in two mutation accumulation line panels that were derived from two wild strains and found background-dependent and fitness-dependent STR mutations. Overall, our results delineate the first large-scale characterization of STR variation in wild *C. elegans* strains and highlight the effects of selection on STR mutations.

## Introduction

Short tandem repeats (STRs) are repetitive elements consisting of 1-6 bp DNA sequence motifs that provide a large source for genetic variation in both inherited and *de novo* mutations (Willems et al. 2016; Fotsing et al. 2019). The predominant mechanism of STR mutations is DNA replication slippage, which often causes expansion or contraction in the number of repeats (Mirkin 2007; Gemayel et al. 2010). Because of their intrinsically unstable nature, STRs have orders of magnitude higher mutation rates than other types of mutations, such as single nucleotide variants (SNVs) and short insertions or deletions (indels) (Lynch 2010; Sun et al. 2012; Willems et al. 2016; Gymrek et al. 2017). The precise mutation rates of STRs are highly variable across different loci and are affected by motif sequences and repeat lengths (Legendre et al. 2007). In humans, STRs are estimated to constitute about 3% of the genome and are associated with dozens of diseases (Mirkin 2007; Hannan 2018; Malik et al. 2021). Emerging studies have also revealed the role of STRs in regulation of gene expression and complex traits in humans and other organisms, which were suggested to facilitate adaptation and accelerate evolution (Fotsing et al. 2019; Jakubosky et al. 2020; Reinar et al. 2021).

The free-living nematode *Caenorhabditis elegans* is a keystone model organism that has been found across the world (Brenner 1974; Kiontke et al. 2011; Andersen et al. 2012; Félix and Duveau 2012; Cook et al. 2017; Crombie et al. 2019; Lee et al. 2021; Crombie et al. 2022). The *C. elegans* Natural Diversity Resource (CeNDR) catalogs and distributes thousands of wild strains, genome sequence data, and genome-wide variation data, including single-nucleotide variants (SNVs) and short indels (Andersen et al. 2012; Cook et al. 2017; Evans et al. 2021). Numerous *C. elegans* population genomics studies and genome-wide association (GWA) studies have leveraged CeNDR resources, such as the genetic variant data across wild strains and the GWA mapping pipeline (Snoek et al. 2020; Evans et al. 2021; Lee et al. 2021; Gilbert et al. 2022; Widmayer et al. 2022; Zhang et al. 2022). However, the natural diversity in *C. elegans* STRs and their impacts on organism-level and molecular traits among wild strains remain unknown because of the lack of STR variation characterization. STRs are challenging to genotype because of their repetitive nature causing errors such as “PCR stutters” (Gymrek 2017). Recent advances provided opportunities to identify genome-wide STR variation accurately in large scales using high-throughput sequencing data (Willems et al. 2017).

In this work, we characterized 31,991 STRs with motif lengths of 1-6 bp in the reference genome of *C. elegans*. We identified natural variation in 27,636 STRs across 540 genetically distinct wild strains and focused on 9,691 polymorphic STRs (pSTRs) with missing calls in less than 10% of all strains. We found enrichment of tri-STRs and hexa-STRs (motif lengths of 3 bp and 6 bp, respectively) in coding (CDS) regions, where STR mutations were likely constrained. Across all pSTRs, we observed more contraction than expansion when we compared wild strains to the reference strain, but the opposite situation could have occurred from the ancestors to descendants. We additionally found that the pSTRs showed similar selective patterns to SNVs, demonstrating that STRs are valuable markers to study population genetics. To further understand STR mutation and evolution in *C. elegans*, we identified 2,956 pSTRs among 174 mutation accumulation lines that were derived from two wild strains using the same STR variant calling pipeline. We found that STR mutation types and rates were affected by genetic background and fitness of ancestors. Our results contribute to the complement of genetic variation characterization in *C. elegans*, develop an efficient STR variant calling pipeline, and provide publicly available resources for future studies.

## Results

### Genome-wide profiling of STR variation in *C. elegans*

To investigate the natural variation of *C. elegans* STRs, we first identified 31,991 reference STRs in the *C. elegans* reference genome (table 1, supplementary table S1). These STRs comprise motif lengths of 1-6 bp and a minimum repeat number of 11, 6, 5, 3.75, 3.4, and 3, respectively for each ascending motif length. The reference STRs were unevenly distributed across the genome (supplementary fig. S1A) with higher density on chromosome arms and tips than centers, suggesting that higher recombination is associated with the increasing incidence of STRs (Rockman and Kruglyak 2009). Mono-STRs (1 bp STRs) that contributed more than half of the reference STRs were also denser on arms and tips than centers, whereas STRs with motif lengths of 2-6 bp distributed differently across the genome (table 1, supplementary fig. S1B).

**Table 1.** The distribution of STRs in *C. elegans*. The numbers and length percentages of polymorphic STRs (reference STRs in parentheses) of different motif lengths in each chromosome and in the whole genome are shown.

| Chromosome | Mono-STR | Di-STR | Tri-STR | Tetra-STR | Penta-STR | Hexa-STR | All STR | Percent of genome (%) |
| --- | --- | --- | --- | --- | --- | --- | --- | --- |
| I | 438<br>(3,094) | 513<br>(1,068) | 360<br>(610) | 171<br>(363) | 37<br>(100) | 148<br>(500) | 1,667<br>(5,735) | 21<br>(91) |
| II | 420<br>(2,662) | 372<br>(737) | 305<br>(515) | 165<br>(349) | 58<br>(135) | 193<br>(593) | 1,513<br>(4,991) | 21<br>(86) |
| III | 393<br>(3,104) | 404<br>(859) | 323<br>(552) | 112<br>(261) | 41<br>(109) | 165<br>(461) | 1,438<br>(5,346) | 21<br>(87) |
| IV | 588<br>(3,162) | 429<br>(867) | 321<br>(608) | 167<br>(354) | 77<br>(168) | 173<br>(429) | 1,755<br>(5,588) | 19<br>(65) |
| V | 443<br>(3,158) | 308<br>(716) | 299<br>(541) | 128<br>(357) | 49<br>(159) | 160<br>(605) | 1,387<br>(5,536) | 13<br>(66) |
| X | 830<br>(2,733) | 512<br>(949) | 261<br>(434) | 114<br>(233) | 67<br>(152) | 145<br>(292) | 1,929<br>(4,793) | 20<br>(52) |
| MtDNA | 1<br>(1) | 0<br>(0) | 0<br>(0) | 1<br>(1) | 0<br>(0) | 0<br>(0) | 2<br>(2) | 21<br>(21) |
| Genome | 3,113<br>(17,914) | 2,538<br>(5,196) | 1,869<br>(3,260) | 858<br>(1,918) | 329<br>(823) | 984<br>(2,880) | 9,691<br>(31,991) | 19<br>(73) |

We examined natural variation in reference STRs across 540 genetically distinct wild *C. elegans* strains (Cook et al. 2017; Evans et al. 2021) and identified 9,691 polymorphic STRs (pSTRs) with missing calls in less than 10% of all strains (table 1, supplementary table S1). The density of pSTRs on arms and tips was not always higher than centers (fig. 1A) likely because DNA slippage, not recombination, is the major source of STR mutations (Kunkel 1993). Poor alignment in hyper-divergent regions (Lee et al. 2021) might also reduce the density of pSTRs in some regions, such as gaps at the left arm of chromosome II and the right arm of chromosome V (fig. 1A). The bases A and T were the most abundant motif sequences in both reference and polymorphic STRs, which is consistent with previous findings in *C. elegans* and many other eukaryotic genomes (Tóth et al. 2000; Denver et al. 2004; Saxena et al. 2019) (fig. 1B, supplementary fig. S3A). We also found that different genomic features were enriched with STRs of different motif lengths (fig. 1C, D, supplementary fig. S3B, C, supplementary table S2). For example, the most prevalent A_n_ and T_n_ mono-STRs were only enriched in introns and intergenic regions (fig. 1D, E, supplementary fig. S3C, D, supplementary table S2). Tri-STRs and hexa-STRs were enriched in CDS regions (fig. 1D, supplementary fig. S3C, supplementary table S2), suggesting purifying selection constrains these STRs to maintain the triplet code (Metzgar et al. 2000). The most enriched tri-STRs in CDS regions were (GGT)_n_ (fig. 1E, supplementary fig. S3D, supplementary table S2), which might correspond to glycine-rich proteins as previously suggested in *C. elegans* (Ying et al. 2016).

**FIG. 1.**
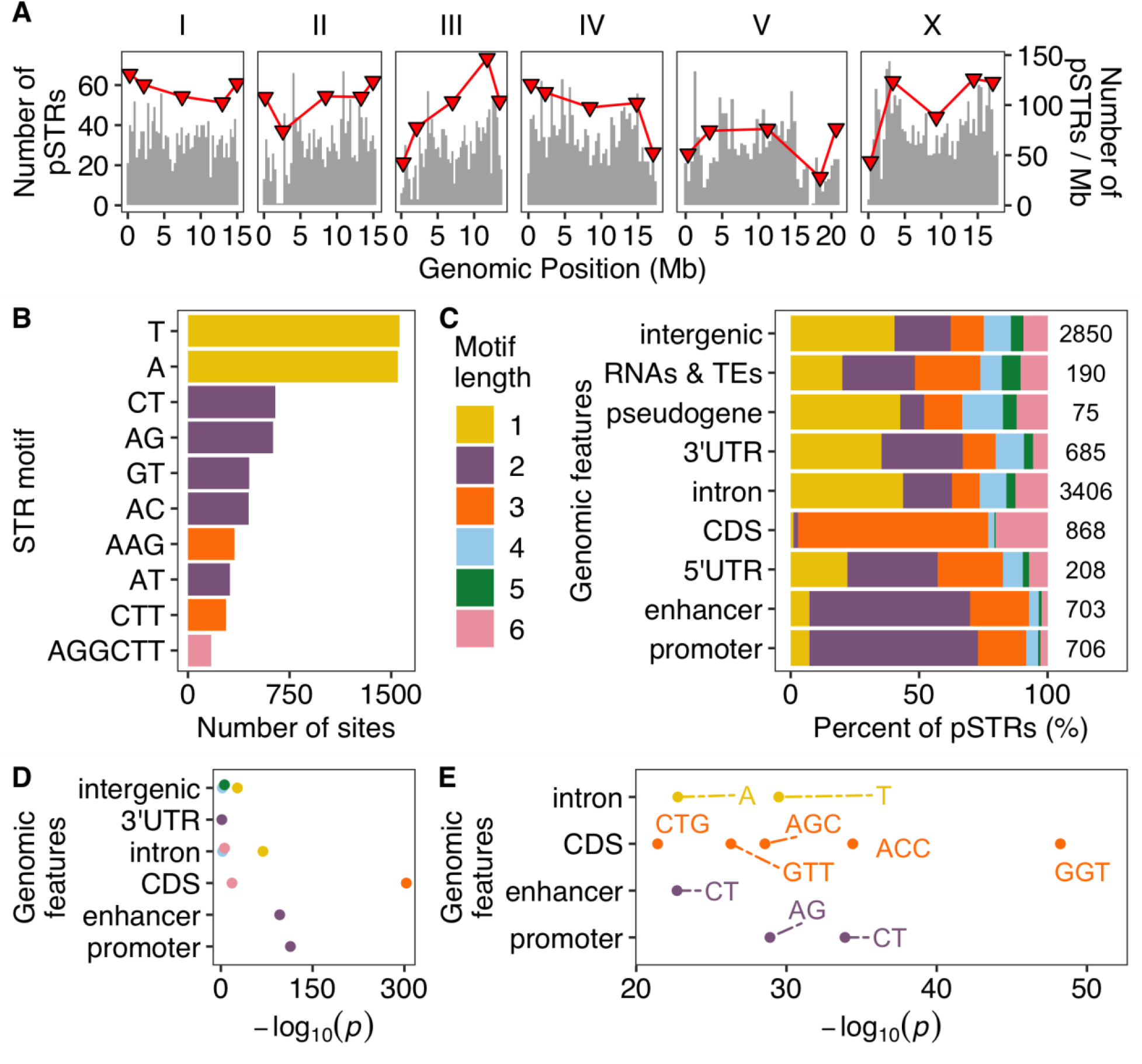
The distribution of polymorphic STRs across *C. elegans. (A)* The distribution of polymorphic STRs (y-axis on the left) in the *C. elegans* genome. Red triangles represent the number of STRs per Mb (y-axis on the right) in different genomic domains (tips, arms, and centers) (Rockman and Kruglyak 2009). *(B)* The top ten most frequent motif sequences in polymorphic STRs are shown on the y-axis, and the number of those sites on the x-axis. *(C)* Percent of polymorphic STRs with different motif lengths in each genomic feature are shown on the x-axis, and different genomic features on the y-axis. The total number of polymorphic STRs in each genomic feature is indicated. *(D)* Enriched STRs with different motif lengths (colored as in (C)) in different genomic features are shown. *(E)* The top 10 most enriched STR motif sequences (labeled) in different genomic features are shown. Statistical significance for enrichment tests (supplementary table S2) was calculated using one-side Fisher’s exact tests and was corrected for multiple comparisons (Bonferroni method).

### Polymorphic STRs are often contracted as compared to the reference genome

STR mutations by DNA slippage are more likely to cause length variation in multiples of the motif lengths (Metzgar et al. 2000; Mirkin 2007) than single nucleotide substitutions. Of alternative alleles among wild *C. elegans* pSTRs, we observed that 30.2%, 35.5%, and 34.3% were insertions, deletions, or substitutions, respectively (fig. 2A). In the same 540 *C. elegans* strains, the proportions of SNVs and indels are 83.3% and 16.7%, respectively. To better understand STR mutations, we computed the expansion and contraction scores (Press et al. 2018) by comparing the longest and/or shortest alternative alleles to the median alleles for each of the 7,506 pSTRs with length variation (fig. 2B). We found significantly higher contraction scores than expansion scores when we compared their absolute values for mono-, tri-, and tetra-STRs (fig. 2C, supplementary table S2). In di-STRs, however, the contraction scores were significantly lower than expansion scores (fig. 2C). Di-STRs stood out as exceptions again in allele frequencies, in which contracted alleles were at significantly lower frequency than expanded alleles (fig. 2D, supplementary table S2). We examined contraction and expansion in STRs with different motif sequences and focused on di-STRs (fig. 2E, supplementary fig. S4). All di-STRs had 36.9% to 38.6% alternative alleles expanded (fig. 2E), except (CG)_n_ di-STRs, which only had 9.6% alternative alleles expanded. This difference might be functionally relevant because no enrichment of (CG)_n_ di-STRs in any genomic features was observed, whereas (AC)_n_, (AG)_n_, (CT)_n_, and (GT)_n_ di-STRs were enriched in promoters and enhancers, and (AT)_n_ di-STRs were enriched in introns. Altogether, we found more STR contraction than expansion among wild *C. elegans*, with the exception of di-STRs.

**FIG. 2.**
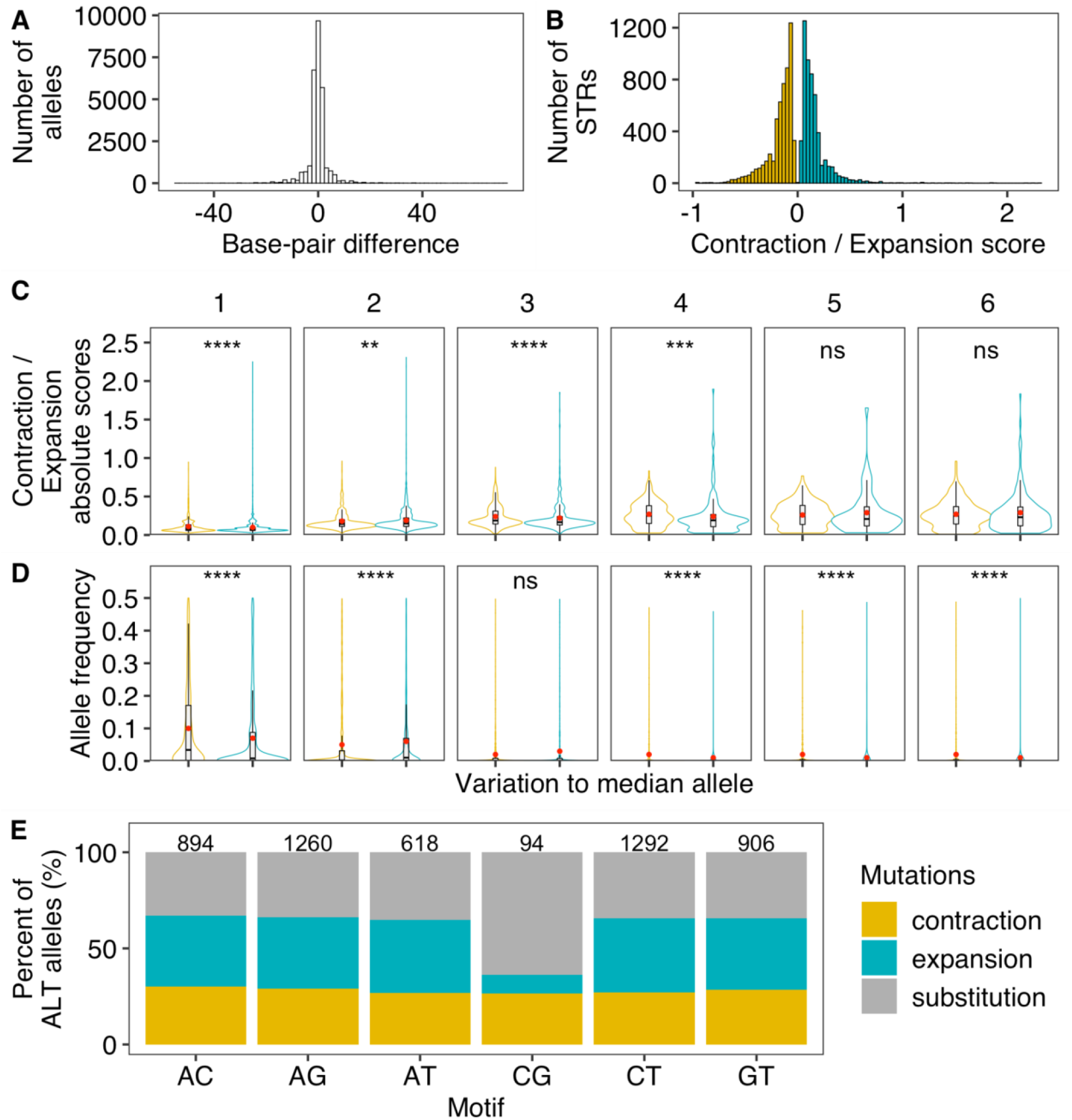
Contraction and expansion of polymorphic STRs. *(A)* The distribution of base-pair differences for polymorphic STR alleles compared to the reference alleles is shown. Positive and negative values on the x-axis indicate allele expansion and contraction, respectively, compared to the reference alleles. *(B)* The distribution of Contraction (in yellow) and Expansion (in blue) scores for each pSTR. Expansion score = [max(STR length) – median(STR length)]/median(STR length); Contraction score = [min(STR length) – median(STR length)]/median(STR length). *(C)* Comparison of the absolute values between Contraction (in yellow) and Expansion (in blue) scores in polymorphic STRs with different motif lengths. *(D)* Comparison of allele frequencies between contracted (in yellow) and expanded alleles compared the median allele length in polymorphic STRs with different motif lengths. Statistical significance was calculated using the two-sided Wilcoxon test and was corrected for multiple comparisons (Bonferroni method). Significance of each comparison (supplementary table S2) is shown above each comparison pair (ns: adjusted *p* > 0.05; *: adjusted *p* ≤ 0.05; **: adjusted *p* ≤ 0.01; ***: adjusted *p* ≤ 0.001; ****: adjusted *p* ≤ 0.0001). *(E)* Percent of alternative alleles showing contraction, expansion, and substitution in di-STRs. The total number of di-STRs with different motif sequences is indicated above each stacked bar.

### STR diversity is correlated with the species-wide selective sweeps

The majority of pSTRs among the 540 wild *C. elegans* strains were multiallelic with a median of three alleles per STR (fig. 3A). Only 4% of pSTRs had a major allele frequency less than 0.5 (fig. 3B), likely because *C. elegans* reproduces primarily by hermaphroditic selfing and recent selective sweeps have reduced diversity across the species (Andersen et al. 2012). The selective sweeps were thought to have purged diversity from the centers of chromosomes I, IV, and V, and the left arm of the X chromosome from the *C. elegans* global population. However, recent sampling efforts of wild *C. elegans* revealed higher genetic diversity in strains from the Hawaiian Islands and other regions in the Pacific Rim, which were hypothesized as the geographic origin of the species (Crombie et al. 2019; Lee et al. 2021; Crombie et al. 2022). We have previously classified wild *C. elegans* into swept and divergent strains based on the proportion of swept haplotypes that were identified using SNVs across the genome (See Materials and Methods) (Crombie et al. 2019; Lee et al. 2021; Zhang et al. 2021). Here, we observed a much higher density of pSTRs with major allele frequencies close to 1 among the 357 swept strains than among all the 540 strains or among the 183 divergent strains (fig. 3B). Within divergent strains, more than 9% of pSTRs had a major allele frequency less than 0.5 (fig. 3B). We also found that divergent strains had a significantly higher percentage of homozygous alternative alleles and heterozygous alleles than swept strains (Wilcoxon test with Bonferroni-corrected *p* = 9.2E-74 and 4.9E-10, respectively) (fig. 3C). Furthermore, principal component analysis (Price et al. 2006) using pSTRs and SNVs showed similar clusters using the 540 strains, which largely correspond to the geographic locations of these strains (fig. 3D, E). The 163 Hawaiian strains, including 157 divergent strains, were mostly separated from the global strains that had experienced the selective sweeps (fig. 3D, E). To further explore the STR diversity in *C. elegans*, we calculated the expected heterozygosity (*H*_*E*_) for each pSTR among all strains, only among swept strains, or only among divergent strains (fig. 3F). The swept strains showed the largest drop of *H*_*E*_ in all the four swept regions (fig. 3F), which is consistent with low levels of genome-wide genetic diversity calculated for these strains using SNVs (Andersen et al. 2012; Crombie et al. 2019; Lee et al. 2021). Divergent strains showed higher diversity across the genome than swept strains and no signatures of selective sweeps (fig. 3F). Altogether, these results suggested that the diversity of STRs in *C. elegans* has been reduced in many strains by the selective sweeps, and divergent strains have retained high levels of STR diversity.

**FIG. 3.**
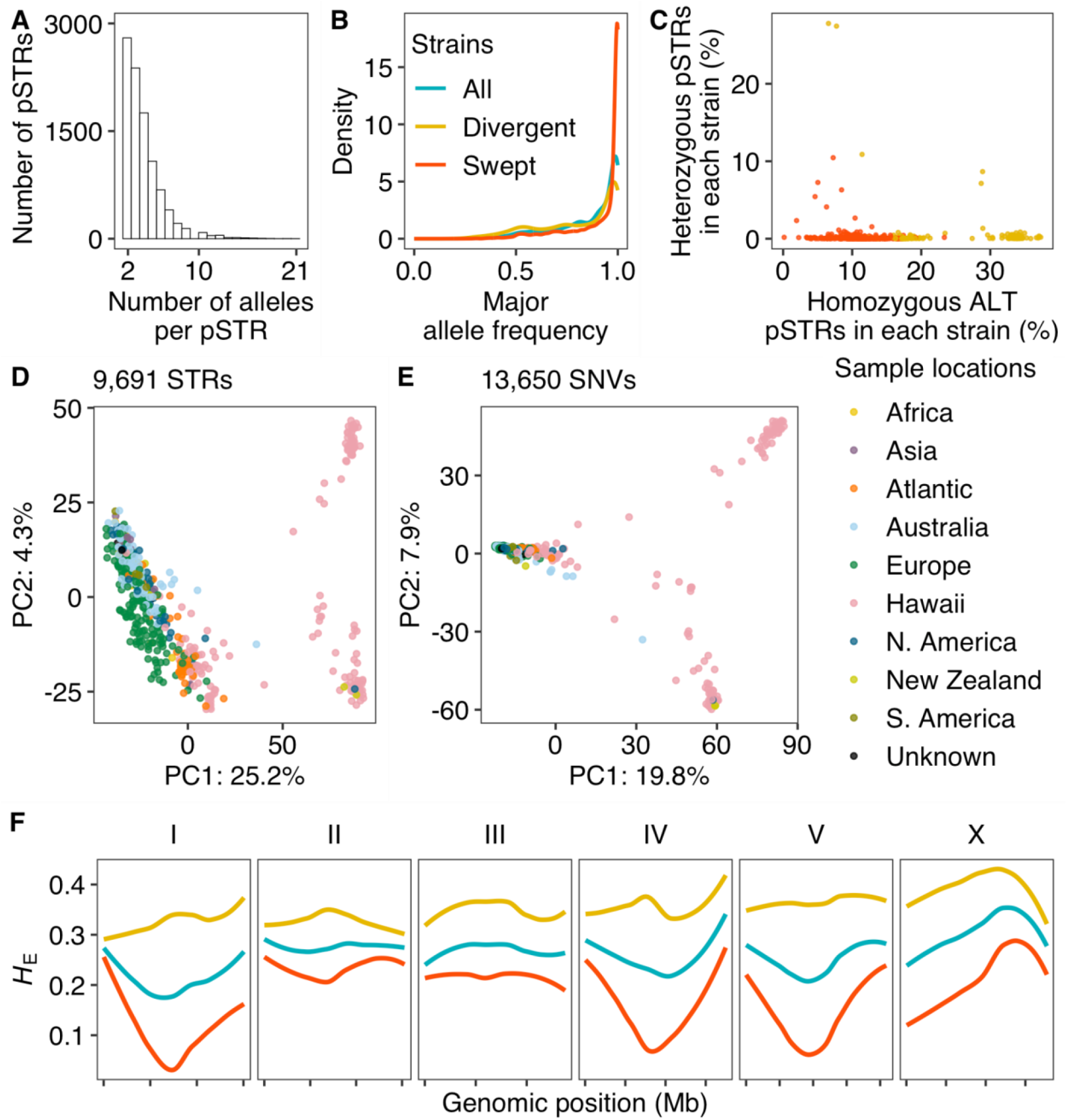
Genetic diversity of *C. elegans* STRs. *(A)* The distribution of allele counts per polymorphic STR. *(B)* Major allele frequencies of all pSTRs for all strains (in blue), divergent strains (in yellow), and swept strains (in red) are shown. *(C)* The percentage of pSTRs with heterozygous alleles is plotted against the percentage of pSTRs with homozygous alternative (ALT) alleles for each of the 540 strains. Divergent and swept strains are colored yellow and red, respectively. *(D, E)* Plots show the top two axes of variation, as determined by principal components analysis (PCA) of the genotype covariances using polymorphic STRs *(D)* and SNVs *(E)*. Each dot represents a strain and is colored by the sampling location. *(F)* Chromosomal expected heterozygosity (*H*_*E*_) of pSTRs is shown as locally regressed lines for all strains (in blue), divergent strains (in yellow), and swept strains (in red). Tick marks on the x-axis denote every 5 Mb.

We further examined pSTR diversity in different genomic features and found that CDS had significantly lower *H*_*E*_ than any other genomic features, indicating reduced pSTR diversity in these regions (supplementary fig. S5A, supplementary table S2). In addition to lower *H*_*E*_, pSTRs in CDS regions also had significantly lower variance in repeat number than most other genomic regions (supplementary fig. S5B, supplementary table S2), suggesting pSTR expansion and contraction might be limited in CDS regions. Increased slippage rates and STR instability were linked to high AT content rather than high GC content (Schlötterer and Tautz 1992; Brandström and Ellegren 2008). We observed the highest GC content among pSTRs in CDS regions (supplementary fig. S5C, supplementary table S2). Altogether, these results suggested that STR diversity was constrained in the conservative CDS regions to maintain proper gene function.

### STR mutation rates in MA lines

In addition to the selective sweeps, other exogenous and endogenous factors might also influence STR diversity in *C. elegans*. For example, because of ample bacterial food and a stable environment (Crombie et al. 2019), Hawaiian strains might have gone through more generations and fewer bottlenecks than European strains, which might have had to enter the dauer diapause stage more frequently to survive starvation and overwinter (Frézal and Félix 2015). Therefore, Hawaiian strains might be able to accumulate more STR and other mutations than European strains. To better understand STR mutation and evolution in *C. elegans*, we examined STR variation in two mutation accumulation (MA) line panels that were derived from two strains, N2 and PB306 (Joyner-Matos et al. 2011; Matsuba et al. 2012; Saxena et al. 2019; Rajaei et al. 2021): 1) N2 MA lines include 67 O1MA lines that were propagated for ∼250 generations, and 38 O2MA lines that were derived from eight selected O1MA lines with high and low fitness and were propagated for an additional ∼150 generations; and 2) PB306 (a wild strain) MA lines include 67 O1MA lines that were propagated for ∼250 generations. We called STR variants using the same methods as for wild strains. We identified 2,956 pSTRs with missing calls in less than 10% of all 172 MA lines and their two ancestors (supplementary fig. S6, supplementary table S3). The pSTRs of MA lines showed similar composition and enrichment features as pSTRs of our 540 wild strains (supplementary fig. S6).

O1MA lines in both MA line panels have undergone passage for about 250 generations with minimal selection (Joyner-Matos et al. 2011; Matsuba et al. 2012; Saxena et al. 2019; Rajaei et al. 2021). To investigate STR mutations in MA lines, we calculated mutation rates for total mutations and three different mutations (deletions, insertions, and substitutions) between the ancestor and each O1MA line (ANC-O1MA) (See Materials and Methods) (fig. 4A-C). We found a significantly lower total mutation rate in O1MA lines derived from the N2 strain than from the PB306 strain (Wilcoxon test with *p* = 0.017) (fig. 4A). Among different types of mutations, N2 O1MA lines showed significantly higher deletion rates but significantly lower substitution rates than PB306 O1MA lines, which were likely driven by mono-STRs (fig. 4B, fig. S7, supplementary table S2). Within each of the two O1MA line panels, we found the highest mutation rates in substitutions (fig. 4B, supplementary table S2). N2 O1MA lines showed significantly higher deletion rates than insertion rates, indicating more contractions than expansions, whereas PB306 O1MA lines showed significantly higher insertion rates than deletion rates (fig. 4B, supplementary table S2). Altogether, these results suggested that genetic variation between the N2 strain and the PB306 strain might affect STR mutation rates and types. Furthermore, we again found that the coding sequence (CDS) had significantly lower mutation rates than all other genomic features, except promoters (fig. 4C, supplementary table S2).

**FIG. 4.**
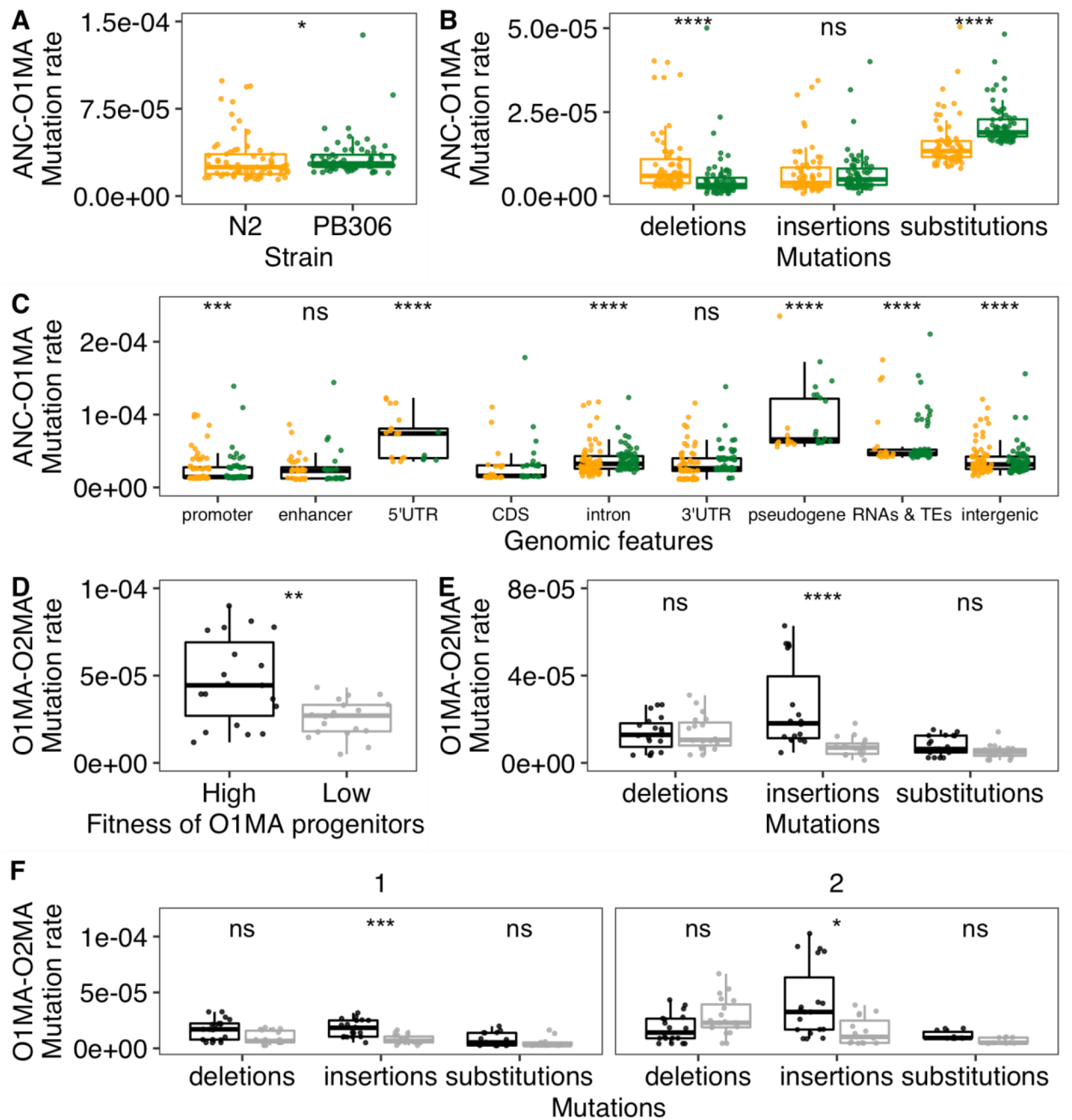
Mutation rates in MA lines. *(A-B)* Comparison of total STR mutation rates *(A)* and STR mutation rates of deletions, insertions, and substitutions *(B)* between O1MA lines derived from N2 (orange) and PB306 (green). *(C)* Comparison of STR mutation rates in CDS regions and other regions using both N2 (orange) and PB306 (green) O1MA lines. *(D-F)* Comparisons of total STR mutation rates *(D)* and STR mutation rates of deletions, insertions, and substitutions using all pSTRs *(E)*, or mono-STRs and di-STRs *(F)* between O2MA lines that were derived from N2 O1MA progenitors with high (black) and low (gray) fitness. Each dot represents the mutation rate between the ancestor strain (ANC) and one of O1MA lines (ANC-O1MA) or between one of the eight N2 O1MA lines and one of its derived O2MA lines (38 in total) (O1MA-O2MA). Statistical significance of difference comparisons (supplementary table S2) was calculated using the two-sided Wilcoxon test and *p*-values were adjusted for multiple comparisons (Bonferroni method). Significance of each comparison is shown above each comparison pair (ns: adjusted *p* > 0.05; **: adjusted *p* ≤ 0.01; ***: adjusted *p* ≤ 0.001; ****: adjusted *p* ≤ 0.0001).

Although minimal selection was maintained during propagation from the N2 ancestor to its O1MA derivatives, lines with consistently high and consistently low fitness at about 250 generations were selected as progenitors for O2MA lines (Matsuba et al. 2012; Saxena et al. 2019). These O2MA lines allowed us to explore how initial fitness (or the initial genomic load of spontaneous deleterious mutations) affects the mutation process of STRs. As for ancestors and O1MA lines, we calculated STR mutation rates between each O1MA line and its O2MA line (O1MA-O2MA) (fig. 4D-F). We found a significantly higher total mutation rate in O2MA lines derived from high fitness O1MA lines than from low fitness O1MA lines (Wilcoxon test with *p* = 0.0056) (fig. 4D). By contrast to ANC-O1MA, the difference in total mutation of O1MA-O2MA was primarily because of insertions rather than substitutions (fig. 4B, E, supplementary table S2). The insertion rates in both mono-STRs and di-STRs were significantly higher in O2MA lines derived from high fitness O1MA lines than from low fitness O1MA lines (fig. 4F, supplementary table S2), whereas deletion rates and substitutions rates using all STRs, mono-STRs, and di-STRs showed no significant differences (fig. 4E, F, supplementary table S2). Altogether, these results suggested which STR mutations might be fitness-dependent, where high-fitness O1MA lines accumulated more STR insertions than low-fitness O1MA lines.

## Discussion

### Natural variation in *C. elegans* STR mutations

STRs have long been recognized as one of the most variable classes of genomic variation. The polymorphisms in few STRs have previously been studied in a limited number of *C. elegans* strains worldwide and in local populations (Sivasundar and Hey 2003; Barrière and Félix 2005; Haber et al. 2005; Barrière and Félix 2007). Here, we characterized the distribution of 31,991 STRs with motif lengths of 1-6 bp in the reference genome of *C. elegans* and identified 9,691 polymorphic STRs across 540 genetically distinct wild strains. We found more STRs on chromosome arms than centers (supplementary fig. S1A), likely because recombination rates were higher on arms than centers (Rockman and Kruglyak 2009). Most pSTRs were multiallelic but had a predominant major allele (fig. 3A, B), which might be caused by the self-fertilizing reproductive mode and deepened by the recent selective sweeps.

As previously demonstrated in other species (Metzgar et al. 2000; Mirkin 2007), length variation caused by deletions or insertions was more common than substitutions among STR mutations in *C. elegans* (fig. 2A). We found significantly more STR contraction than expansion (fig. 2B, C) when we compared wild strain genomes to the reference genome. The reference strain N2 was isolated from Bristol, England and was identified as a swept strain (Andersen et al. 2012). To understand the evolution of STRs in *C. elegans*, a more informative comparison might come from choosing a reference from strains that avoided selective sweeps and was isolated from regions nearby the species origins. For example, the Hawaiian strain XZ1516, which likely carries the most ancestral genotypes (Crombie et al. 2019; Ma et al. 2021), has contraction in 66% of the 2,400 pSTRs that showed length variation to the reference STRs. Therefore, STR expansion rather than contraction likely occurred from ancestors to descendants in *C. elegans* if we consider strains that might reflect the ancestral origin of the species.

### Polymorphic STRs reflect species evolutionary history

The differences in SNV diversity across the genomes of wild strains revealed the species-wide selective sweeps and potential geographical origins of *C. elegans* (Andersen et al. 2012; Crombie et al. 2019; Lee et al. 2021; Crombie et al. 2022). Our results in STR diversity across the *C. elegans* genome of the 540 wild strains agreed with previous discoveries using SNVs. STR diversity across the genome showed signatures of selective sweeps among the 357 swept strains (fig. 3F) in similar genomic regions as previous results (Andersen et al. 2012). We found higher STR diversity in divergent strains than swept strains (fig. 3B, C). The divergence in STRs across wild strains corresponded to their geographic locations as revealed by SNVs (fig. 3D, E). Altogether, these results suggest natural variation in STRs reflect the evolutionary history of *C. elegans*. Because of the higher mutation rates of STRs than SNVs, exploring STR polymorphisms could further help to better resolve demography and short-scale genealogy in population genetic studies.

### The impacts of selection on STR variation

The species-wide selective sweeps might have had significant influences on the STR diversity that we observed in the wild *C. elegans* strains (fig. 3). Additionally, purifying selection might have constrained motif lengths and mutations of STRs in CDS regions to maintain proper functions in wild strains (fig. 1D, E, supplementary fig. S3C, D, supplementary fig. S5). We also observed constrained STRs in CDS regions of MA lines (fig. 4C, supplementary fig. S6C, D), which in principle mostly experienced relaxed selection, indicating strong deleterious effects of STR variation on CDS functions. We found the highest mutation rates in substitutions of ANC-O1MA (fig. 4B) and in insertions of O1MA-O2MA (fig. 4E), which might be related to their different mutation loads in the progenitors, because the growth environment from the ancestor to O1MA and from O1MA to O2MA was essentially identical (Matsuba et al. 2012; Saxena et al. 2019). It would be interesting to investigate the mutation pattern in a narrow time range, for example, each 50 generations, to examine if mutation types and rates are associated with the load of mutations accumulated in the background during the spontaneous mutational process.

Among O2MA lines, we also found fitness-dependent STR mutations (fig. 4D-F). O2MA lines derived from high fitness O1MA lines showed significantly higher insertion rates than those derived from low fitness O1MA lines (fig. 4E, F). The original study found the short indel mutation rate was significantly greater in the high fitness lines than in the low fitness lines (Saxena et al. 2019). The authors proposed that high fitness lines might have higher tolerance than low fitness lines to harbor more indels because of synergistic epistasis (Saxena et al. 2019), which might also explain the fitness-dependent STR mutation that we observed here. Expansion could decrease the stability of STRs and has been widely associated with human disease and trait defects (Mirkin 2007; Sureshkumar et al. 2009). Assuming expanded STRs are more likely to have deleterious effects on fitness than contracted STRs, high-fitness MA lines might be able to accumulate more expanded STRs than low fitness lines before being removed by selection. Future effects should measure the fitness of O2MA lines and examine the correlation between STR mutation rates and fitness.

## Materials and Methods

### *C. elegans* genotype data

We obtained the reference genome of *C. elegans* from WormBase (WS276) (Harris et al. 2020) and alignment of whole-genome sequence data in the BAM format of 540 wild *C. elegans* strains from CeNDR (20210121 release) (Andersen et al. 2012; Cook et al. 2017; Evans et al. 2021). These BAM files were generated using *BWA (Li and Durbin 2009)* incorporated in the pipeline *alignment-nf* (https://github.com/AndersenLab/alignment-nf) (Cook et al. 2017). We also acquired the hard-filtered isotype variant call format (VCF) file (CeNDR 20210121 release) for SNVs among the 540 wild *C. elegans* strains (Cook et al. 2017).

### STR variant calling

We built a STR reference from the *C. elegans* reference genome using *Tandem Repeats Finder (Benson 1999)* and the STR reference construction framework described in *HipSTR-references* (https://github.com/HipSTR-Tool/HipSTR-references) (Willems et al. 2017). Then, we called STR variants using BAM files of the 540 strains, the STR reference, and *HipSTR* (v0.6.2) in the *de novo* stutter estimation mode (Willems et al. 2017). We filtered the VCF of *HipSTR* output using the script *filter_vcf*.*py* as recommended in *HipSTR* to have high-quality calls. In total, we found variation in 27,636 STRs among the 540 strains. We further filtered STR variants with equal or more than 10% missing data across all strains using *BCFtools* (v.1.9) (Li 2011) and came to 9,691 polymorphic STRs, which we used in downstream analyses.

### STR annotation and effect prediction

We determined genomic regions of reference STRs according to the general feature format (GFF3) file from WormBase (WS276) (Harris et al. 2020) and prediction of promoters and enhancers (Jänes et al. 2018). STRs with multiple annotated features were assigned to a single feature using the following priority: CDS > 5’UTR > 3’UTR > promoter > enhancer > intron > RNAs & TEs > intergenic regions. We further predicted the consequence of polymorphic STR variants using the *csq* function of *BCFtools* (v.1.12) (Li 2011) incorporated in the pipeline *annotation-nf* (https://github.com/AndersenLab/annotation-nf).

### Expansion and contraction

For each polymorphic STR with expanded and/or contracted alternative alleles, we calculated the Expansion score = [max(STR length) – median(STR length)]/median(STR length) (Press et al. 2018) and/or the Contraction score = [min(STR length) – median(STR length)]/median(STR length).

### Classification of swept and divergent strains

We acquired the sweep haplotype summary data of the 540 wild *C. elegans* strains from CeNDR (20210121 release) (Cook et al. 2017). We defined strains with greater than or equal to 30% of swept haplotype in any of the four chromosomes (I, IV, V, and X) as swept strains. Other strains were defined as divergent strains.

### Principal components analysis (PCA)

For STRs, because only eight polymorphic STRs have no missing data for all 540 strains, we imputed the genotype of the 9,691 polymorphic STRs for strains with missing data. For strains with homozygous alleles (*e*.*g*., “0|0”, “1|1”, “2|2”), a single character (*e*.*g*., “0”, “1”, “2”), was used to represent the genotype. For strains with heterozygous alleles (*e*.*g*., “0|1”, “1|2”, “3|2”), we treated the genotypes as numeric values and chose the smaller one as the genotype (*e*.*g*., “0”, “1”, “2”). Then we imputed missing genotypes using the R package *missMDA* (v1.18) (Josse and Husson 2016). For SNVs, we used the hard-filtered isotype VCF (CeNDR 20210121 release) and used *BCFtools* (Li 2011) to filter SNVs that had any missing genotype calls and those that were below the 5% minor allele frequency. We used *PLINK* v1.9 (Purcell et al. 2007; Chang et al. 2015) to prune the SNVs to 13,650 markers with a linkage disequilibrium threshold of *r*^*2*^ < 0.8. Then, we used the generic function *prcomp()* in R (Core Team and Others 2013) to perform principal components analysis for both STRs and SNVs.

### STR diversity

We calculated expected heterozygosity (*H*_*E*_) (Nei 1973) for STR diversity using the following equation:

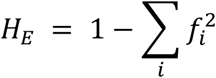

where the *f_i_* denotes the allele frequency of the *i*th allele for a specific STR.

### STR variants in mutation accumulation (MA) lines

We obtained whole-genome sequence data in the FASTQ format of 174 MA lines, including N2 MA lines: the N2 ancestor, 67 O1MA lines, and 38 O2MA lines; PB306 MA lines: the PB306 ancestor and 67 O1MA lines (NCBI Short Read Archive projects PRJNA395568, PRJNA429972, and PRJNA665851) (Saxena et al. 2019; Rajaei et al. 2021). We used the pipelines *trim-fq-nf* (https://github.com/AndersenLab/trim-fq-nf) and *alignment-nf* to trim raw FASTQ files and generate BAM files for each line, respectively. We called STR variants for the 174 lines as described above and identified 2,956 pSTRs with missing calls in less than 10% of all strains.

### Mutation rate of polymorphic STRs in MA lines

We calculated the STR mutation rate in MA lines using their 2,956 pSTRs. For each O1MA line and the ancestor, we selected STR sites with data in both lines. Then, we compared the two alleles of each STR in the O1MA line to the two alleles in the ancestor, respectively, to identify insertion, deletion, substitution, or no mutation. We also obtained the number of generations between the O1MA line and the ancestor from the original studies (Saxena et al. 2019; Rajaei et al. 2021). The mutation rate (per-allele, per-STR, per-generation) *µ* for each type of mutation was calculated as *m/2nt* where *m* is the number of the mutation, *n* is the total number of STR sites between the two lines, and *t* is the number of generations. We calculated the mutation rate of the three different mutations for each N2 O2MA line to its ancestral N2 O1MA line using the same method.

### Statistical analysis

Statistical significance of difference comparisons was calculated using the Wilcoxon test and *p*-values were adjusted for multiple comparisons (Bonferroni method) using the *compare_means()* function in the R package *ggpubr* (v0.2.4) (https://github.com/kassambara/ggpubr/). Enrichment analyses were performed using the one-sided Fisher’s exact test and were corrected for multiple comparisons (Bonferroni method).

## Acknowledgments

We would like to thank Timothy A. Crombie and Ryan McKeown for helpful comments on the manuscript. We would also like to thank WormBase because without it these analyses would not have been possible. G.Z. is supported by the NSF-Simons Center for Quantitative Biology at Northwestern University (awards Simons Foundation/SFARI 597491-RWC and the National Science Foundation 1764421). Y.W. was supported as a joint PhD student by China Scholarship Council (No. 201706910052). E.C.A. is supported by a grant from the National Institutes of Health R01 DK115690. The *C. elegans* Natural Diversity Resource is supported by a National Science Foundation Living Collections Award to E.C.A. (1930382).

## Author contributions

E.C.A. conceived of and designed the study. G.Z. and Y.W. analyzed the data. G.Z., Y.W., and E.C.A. wrote the manuscript.

## Competing Interests

The authors declare no competing interests.

## Data availability

The STR variant calling pipeline can be found at https://github.com/AndersenLab/wi-STRs. The datasets and code for generating all figures can be found at https://github.com/AndersenLab/WI-Ce-STRs.

## Supplementary material

### Supplementary Figures

**Fig. S1.**
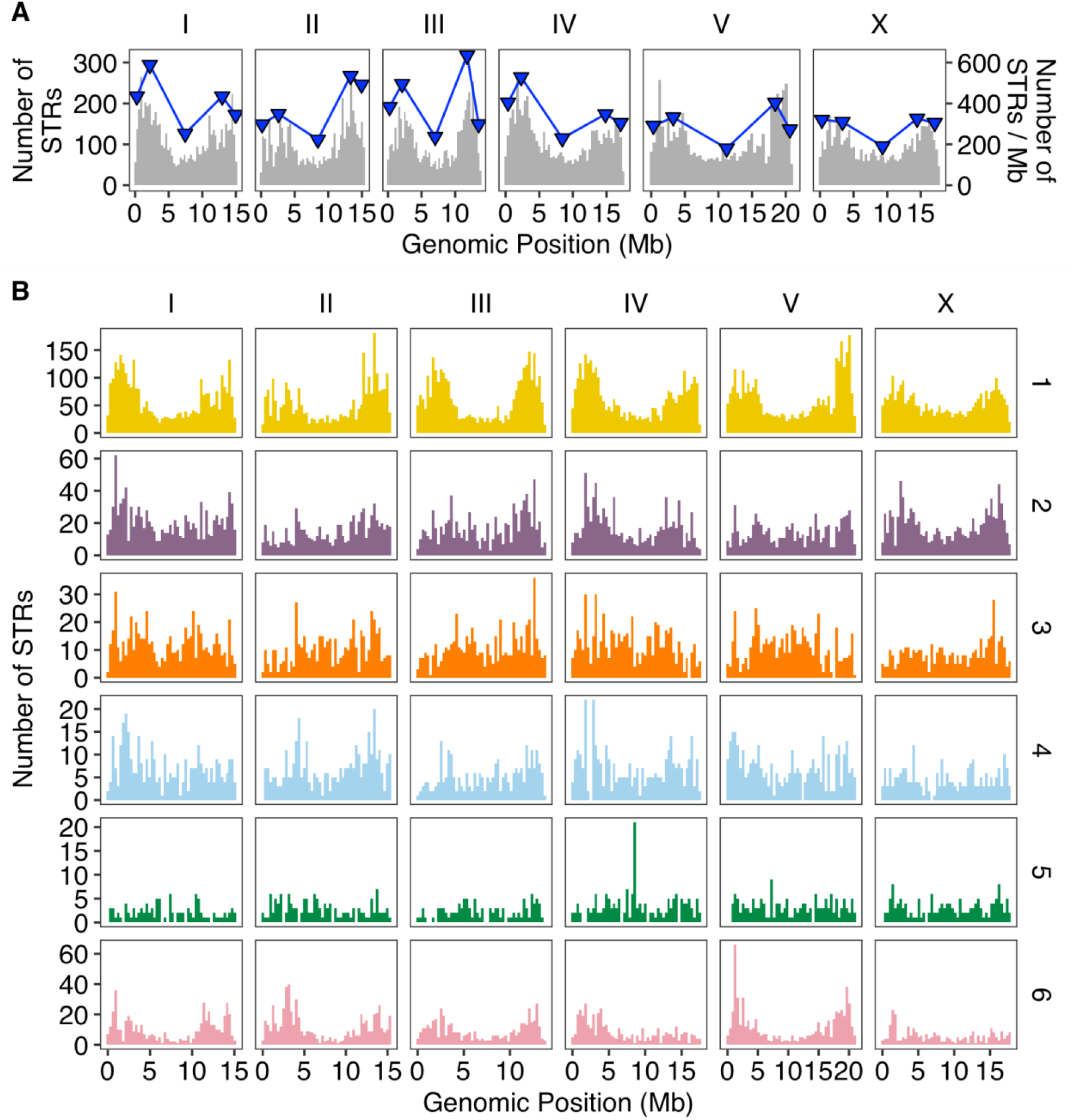
The distribution of reference STRs across *C. elegans. (A)* The distribution of reference STRs in the *C. elegans* genome. Blue triangles represent the number of STRs per Mb in different genomic domains (tips, arms, and centers) (Rockman and Kruglyak 2009). *(B)* The distribution of reference STRs with different motif lengths in the *C. elegans* genome.

**Fig. S2.**
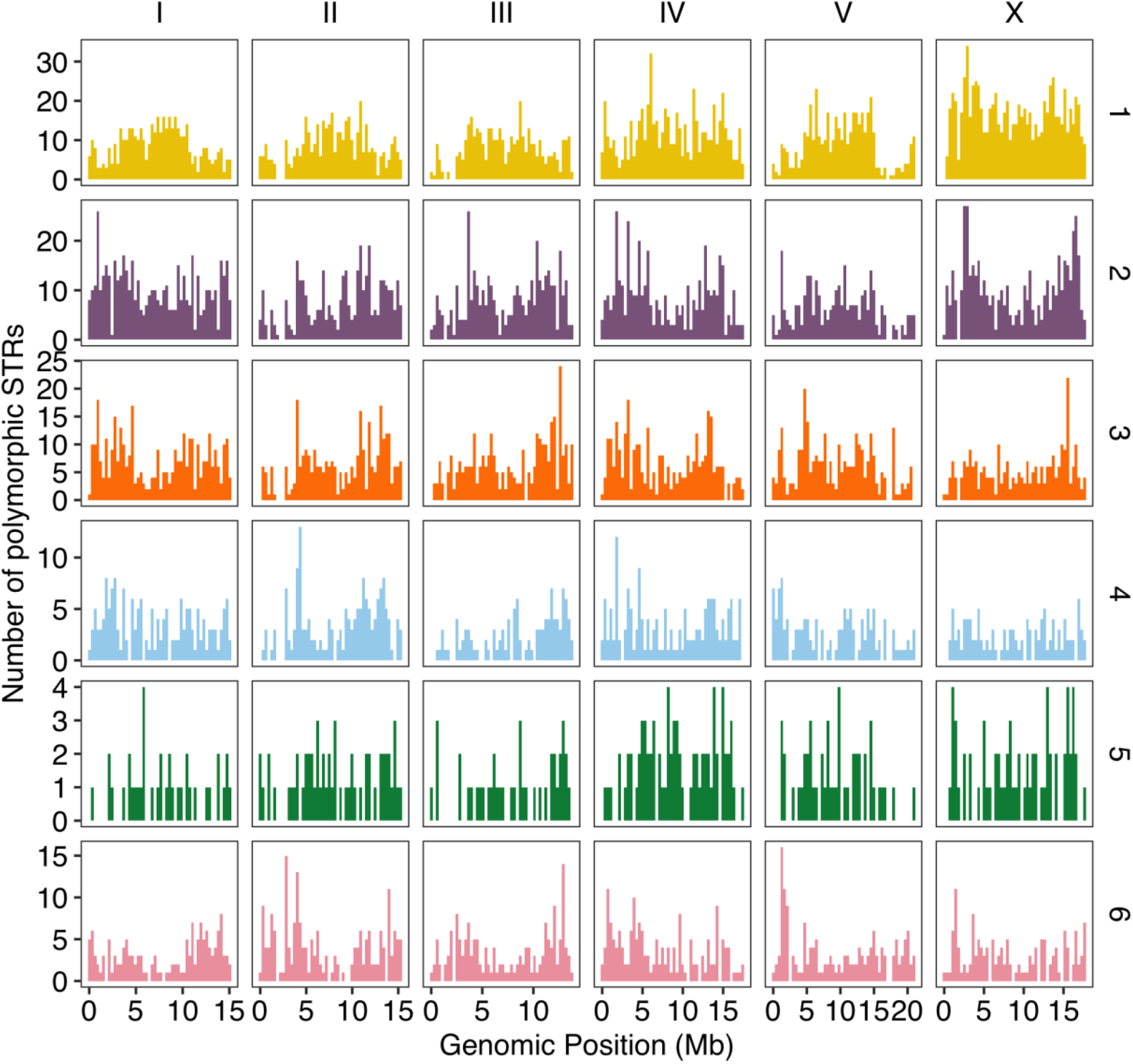
The distribution of polymorphic STRs with different motif lengths in the *C. elegans* genome.

**Fig. S3.**
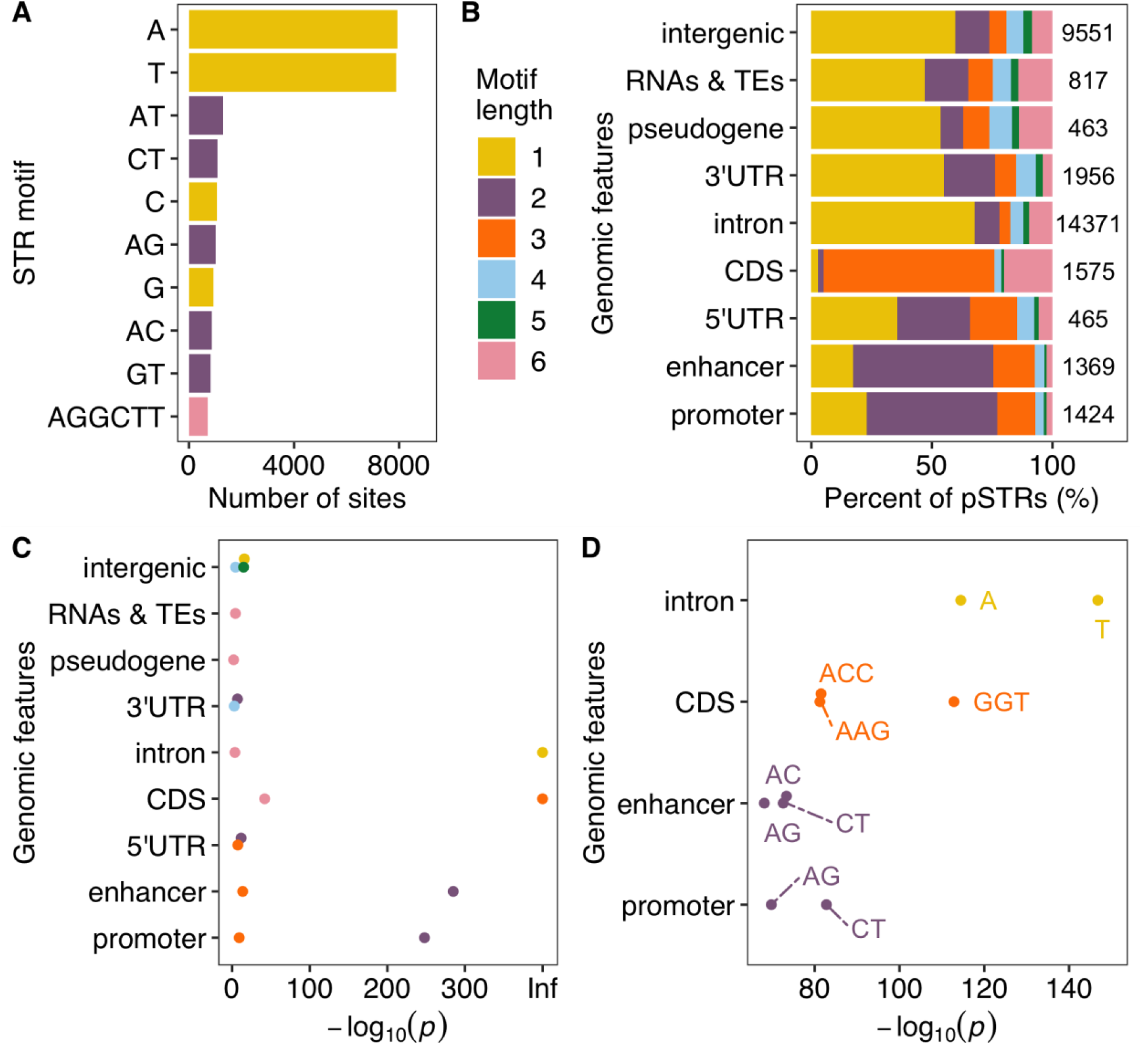
Motifs and genomic features of reference STRs in *C. elegans. (A)* The top ten most frequent motif sequences in reference STRs are shown on the y-axis, and the number of those sits on the x-axis. *(B)* Percent of reference STRs with different motif lengths in each genomic feature. The total number of reference STRs in each genomic feature is indicated. *(C)* Enriched STRs with different motif lengths (colored as in (B)) in different genomic features are shown. *(D)* The top 10 most enriched STR motif sequences (labeled) in different genomic features are shown. Statistical significance (supplementary table S2) for enrichment tests was calculated using the one-side Fisher’s exact test and was corrected for multiple comparisons (Bonferroni method).

**Fig. S4.**
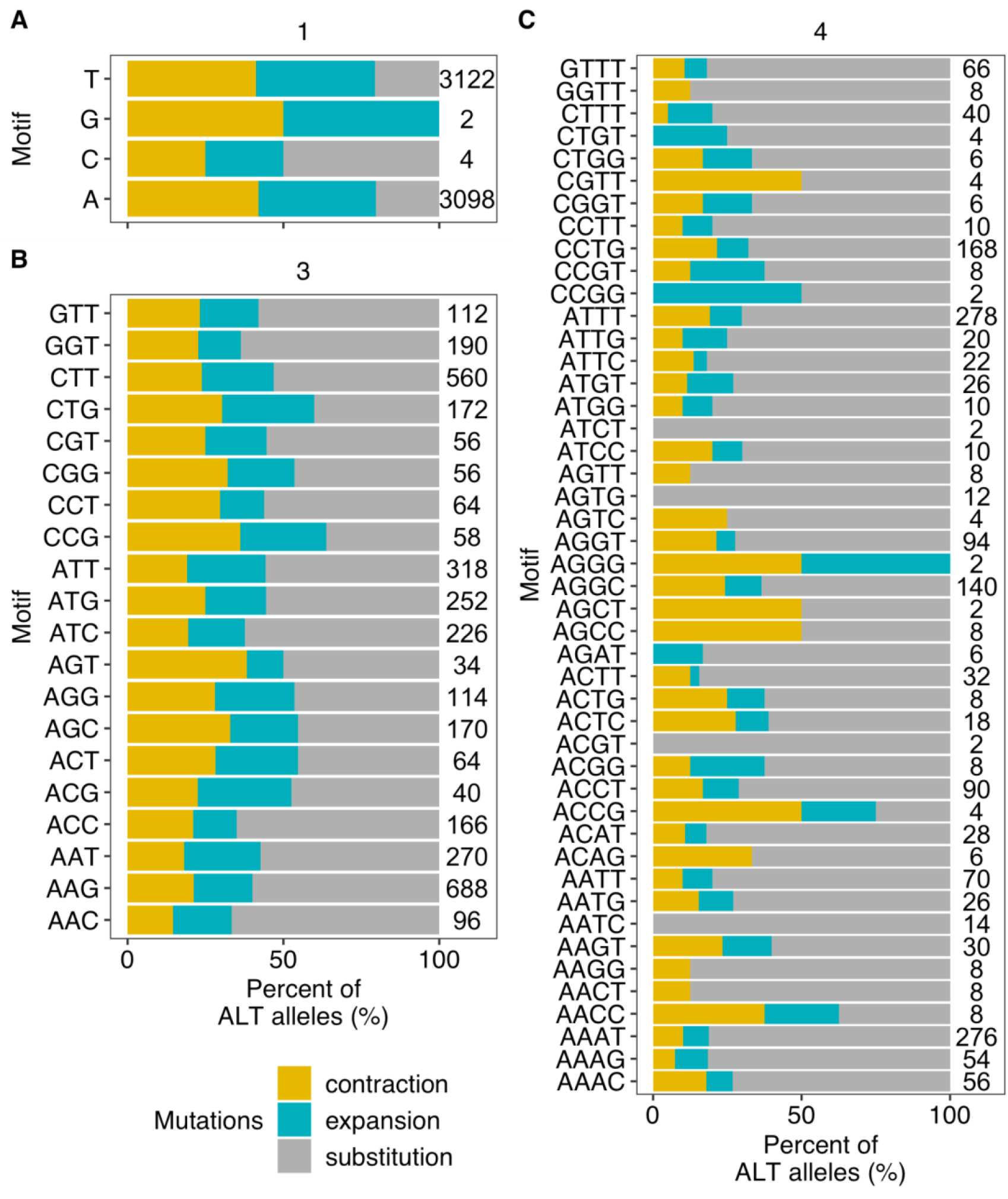
Percent of alternative alleles showing contraction, expansion, or substitution in mono-STRs *(A)*, tri-STRs *(B)*, and tetra-STRs *(C)*. The total number of STRs with different motifs is indicated on the right.

**Fig. S5.**
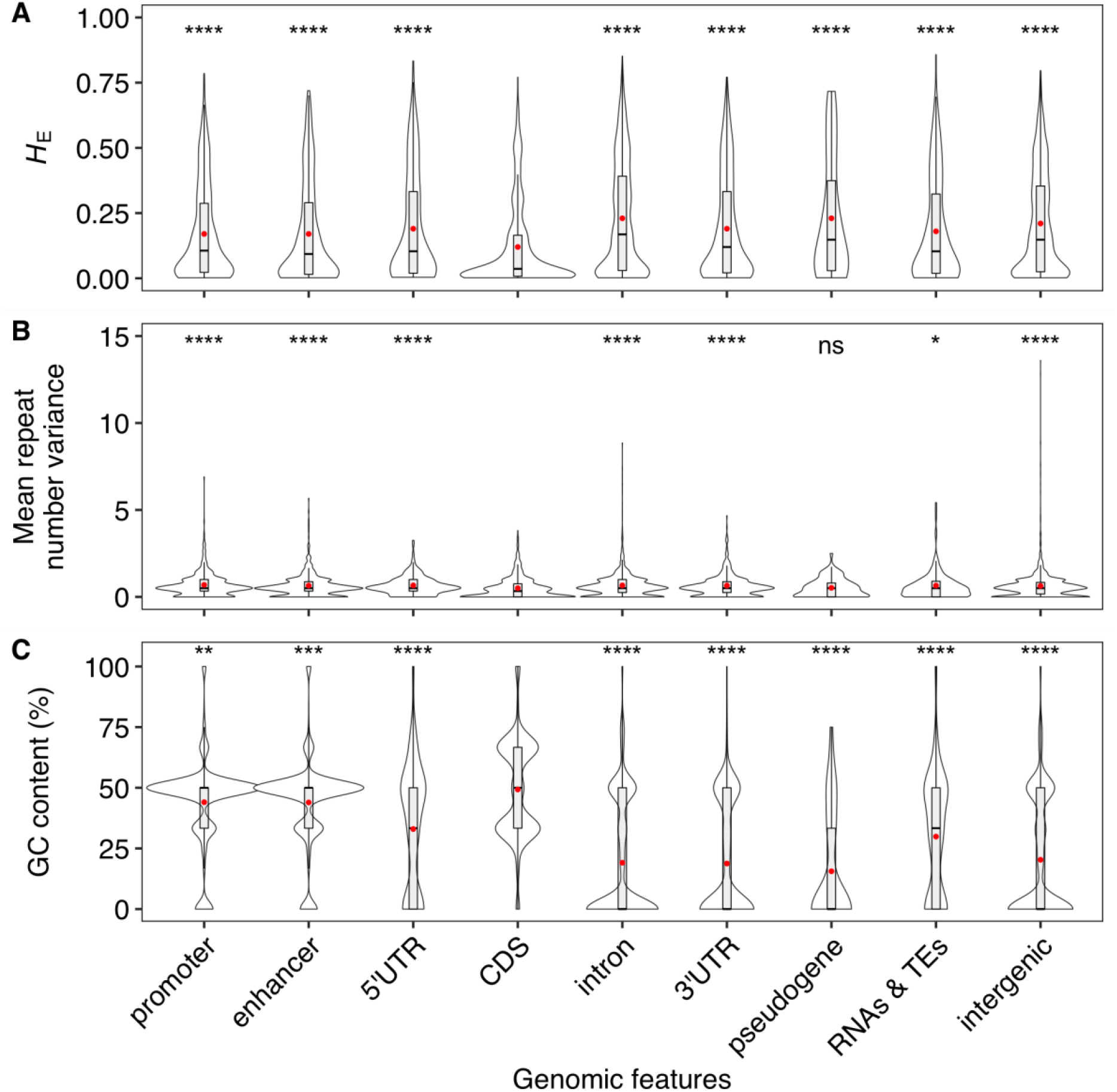
Mutations of STRs in CDS regions are constrained. Comparisons of expected heterozygosity (*H*_E_) *(A)*, mean repeat number variance of each STRs *(B)*, and motif GC content *(C)* between polymorphic STRs in CDS regions and other regions. Red dots indicate mean values of each estimate in each region. Statistical significance (supplementary table S2) was calculated using the two-sided Wilcoxon test and was corrected for multiple comparisons (Bonferroni method). Significance of each comparison is shown above (ns: adjusted *p* > 0.05; *: adjusted *p* ≤ 0.05; **: adjusted *p* ≤ 0.01; ***: adjusted *p* ≤ 0.001; ****: adjusted *p* ≤ 0.0001).

**Fig. S6.**
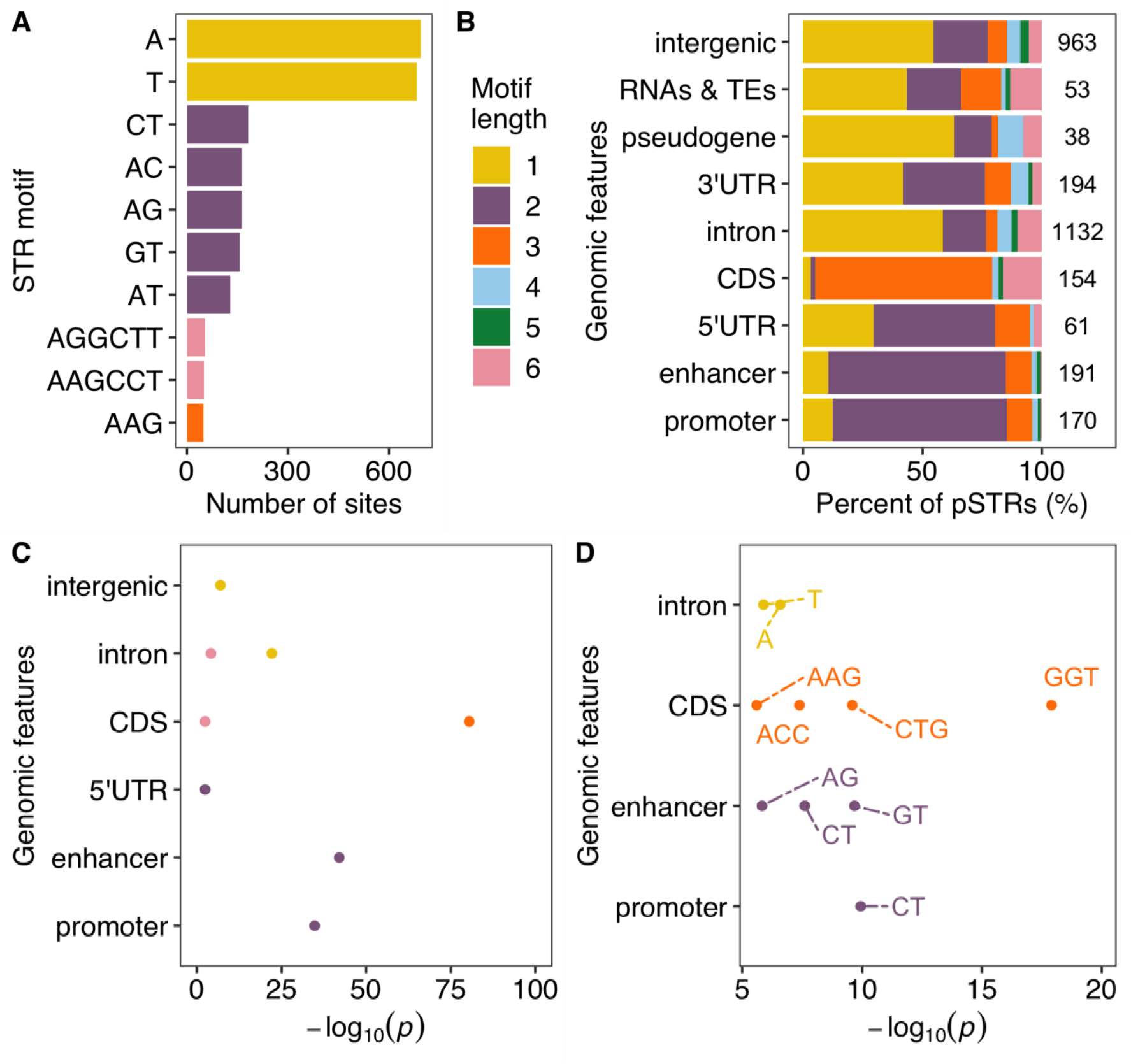
Motifs and genomic features of polymorphic STRs in MA lines. *(A)* The top ten most frequent motif sequences in polymorphic STRs. *(B)* Percent of polymorphic STRs with different motif lengths in each genomic feature. The total number of polymorphic STRs in each genomic feature are indicated. *(C)* Enriched STRs with different motif lengths (colored as in (B)) in different genomic features are shown. *(D)* Enriched STR motif sequences (labeled) in different genomic features are shown. Statistical significance (supplementary table S2) for enrichment tests was calculated using the one-side Fisher’s exact test and was corrected for multiple comparisons (Bonferroni method).

**Fig. S7.**
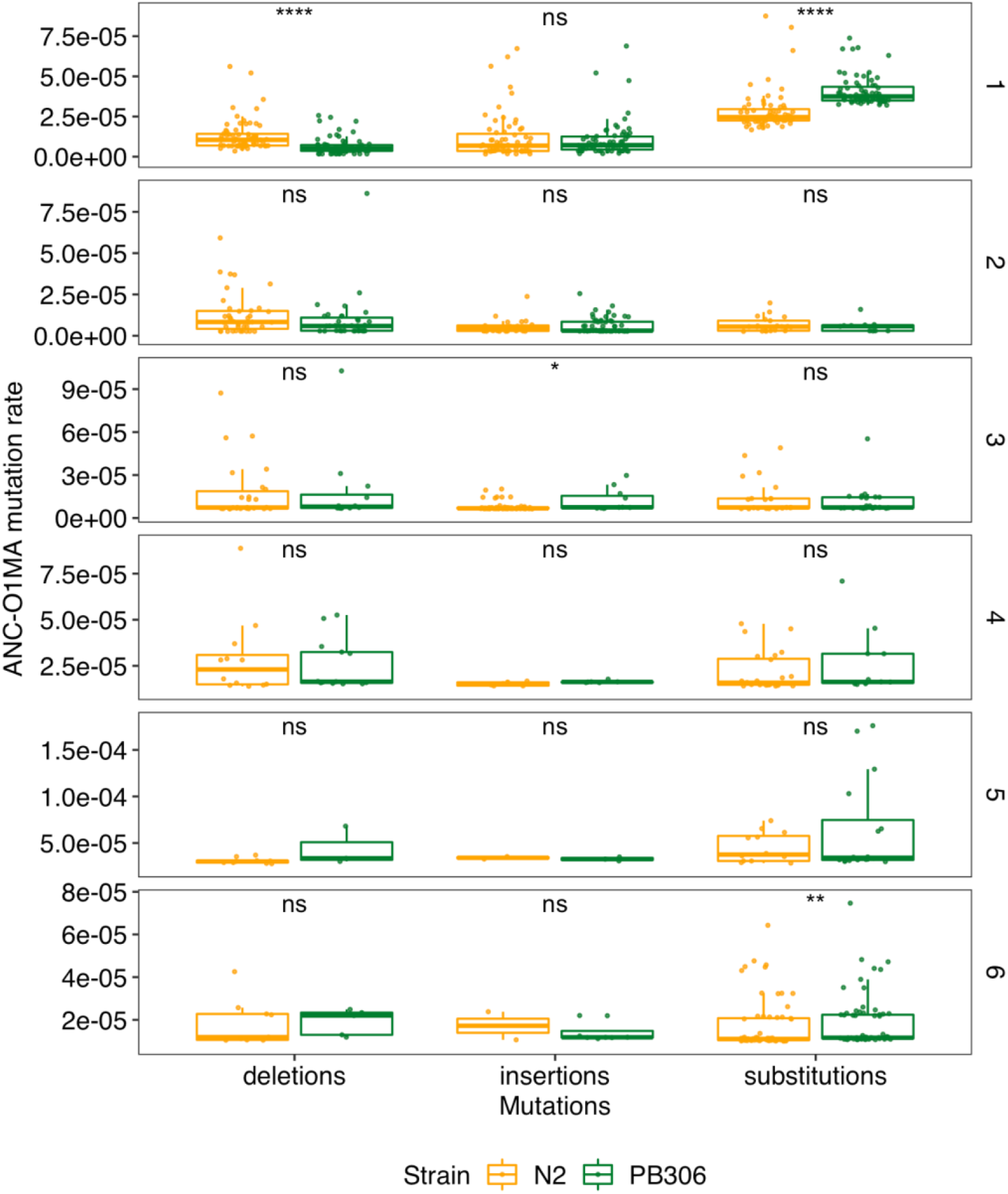
Comparison of STR mutation rates of deletions, insertions, and substitutions between O1MA lines derived from N2 (orange) and PB306 (green) using pSTRs of different motif lengths. Each dot represents the mutation rate between the ancestor strain (ANC) and one of O1MA lines (ANC-O1MA). Statistical significance (supplementary table S2) of difference comparisons were calculated using the two-sided Wilcoxon test and *p*-values were adjusted for multiple comparisons (Bonferroni method). Significance of each comparison is shown above each comparison pair (ns: adjusted *p* > 0.05; **: adjusted *p* ≤ 0.01; ***: adjusted *p* ≤ 0.001; ****: adjusted *p* ≤ 0.0001).

### Supplementary Tables

**Table S1**

Reference STRs, polymorphic STRs, and pSTR expansion/contraction scores among wild *C. elegans* strains

**Table S2**

Exact adjusted *p* values in FIGs. 1D-E, 2C-D, 4B-C, E-F, S3C-D, S5, S6C-D, S7

**Table S3**

Polymorphic STRs found in the MA lines.

## Notes

### Competing Interest Statement

The authors have declared no competing interest.

